# EEG-based clusters differentiate psychological distress, sleep quality and cognitive function in adolescents

**DOI:** 10.1101/2021.10.14.464347

**Authors:** Owen Forbes, Paul E. Schwenn, Paul Pao-Yen Wu, Edgar Santos-Fernandez, Hong-Bo Xie, Jim Lagopoulos, Larisa T. McLoughlin, Dashiell D. Sacks, Kerrie Mengersen, Daniel F. Hermens

## Abstract

1

**Introduction:** To better understand the relationships between brain activity, cognitive function and mental health risk in adolescence there is value in identifying data-driven subgroups based on measurements of brain activity and function, and then comparing cognition and mental health symptoms between such subgroups.

**Methods:** Here we implement a multi-stage analysis pipeline to identify data-driven clusters of 12-year-olds (M = 12.64, SD = 0.32) based on frequency characteristics calculated from resting state, eyes-closed electroencephalography (EEG) recordings. EEG data was collected from 59 individuals as part of their baseline assessment in the Longitudinal Adolescent Brain Study (LABS) being undertaken in Queensland, Australia. Applying multiple unsupervised clustering algorithms to these EEG features, we identified well-separated subgroups of individuals. To study patterns of difference in cognitive function and mental health symptoms between core clusters, we applied Bayesian regression models to probabilistically identify differences in these measures between clusters.

**Results:** We identified 5 core clusters which were associated with distinct subtypes of resting state EEG frequency content. EEG features that were influential in differentiating clusters included Individual Alpha Frequency, relative power in 4 Hz bands up to 16 Hz, and 95% Spectral Edge Frequency. Bayesian models demonstrated substantial differences in psychological distress, sleep quality and cognitive function between these clusters. By examining associations between neurophysiology and health measures across clusters, we have identified preliminary risk and protective profiles linked to EEG characteristics.

**Conclusion:** In this work we have developed a flexible and scaleable pipeline to identify subgroups of individuals in early adolescence on the basis of resting state EEG activity. These findings provide new clues about neurophysiological subgroups of adolescents in the general population, and associated patterns of health and cognition that are not observed at the whole group level. This approach offers potential utility in clinical risk prediction for mental and cognitive health outcomes throughout adolescent development.

## 2 Introduction

Mental health issues are a significant and growing problem in youth populations, and confer significant burden on public health, communities and individual wellbeing (1; 2). Adolescence is a period in which many mental health problems are thought to emerge, as the brain is rapidly developing and changing (3; 4). It is also a crucial period in which to identify emerging psychopathology, to anticipate risk trajectories and enable early intervention (2). Given the high burden of mental health problems in adolescents along with the overlapping risk across disorder diagnoses in this age range, there is urgent need for novel data-driven methods to support improved diagnoses, risk prediction and early intervention for psychopathology and cognitive development in adolescence.

In this work we report on a flexible analysis pipeline that enables identification of subgroups of individuals based on frequency features of electroencephalography (EEG) data, and allows comparison of differences in neurophysiology and health measures between these subgroups. Our aim was to identify empirical, data-driven patterns relating brain activity to cognitive function and mental health outcomes that are more robust and generalisable than the findings available through a case-control approach. Traditional approaches to characterising individuals based on brain measurements typically rely on a case-control approach, which separates individuals into groups on the basis of a clinical diagnosis, contrasted with ‘healthy controls’ without any diagnosis (5). There are several shortcomings to the case-control approach, including that it is bound to find patterns of difference between groups categorised by traditional diagnostic structures based on clinician judgement. As a result case-control studies have limited ability to generalise and capture information relating to psychopathology and cognitive function issues occurring on a continuum, and are limited to either contradicting or strengthening existing assumptions about these categorical differences. In this framework, differences in brain characteristics of interest between cases and controls are also likely to be confounded by many extraneous factors (5). Recent studies have suggested that EEG studies of mental health should strive to be more independent of clinical diagnostic categories and examine mental health symptoms from a data-driven neurophysiology perspective (6; 7).

Previous research has demonstrated that features calculated from EEG data are associated with mental health and cognitive function outcomes across multiple life stages (8; 6). Particularly in the frequency domain, a large variety of studies have demonstrated these sorts of associations. For example, the ratio of theta to beta activity has been used to support diagnosis and prognosis in individuals with attention deficit hyperactivity disorder (ADHD) (9). Resting state theta power has been associated with cognitive performance in children with ADHD (10). Various EEG-based biomarkers have been associated with treatment response for depression in adults (11). EEG signal complexity or entropy has been shown to be associated with neurological conditions including epilepsy and autism spectrum disorders (12; 13). These are just a few of the many studies which have identified differences between controls and individuals with a particular diagnosis, based on specific features calculated from EEG data. By studying novel combinations of summary features in a data-driven framework, we can develop unique EEG-based characterisations of individuals and investigate associations between distinct EEG profiles and health outcomes of interest.

Mental disorder risk in childhood and adolescence is understood to be pluripotential, with non-specific risk overlapping across a number of categorical diagnostic outcomes (14; 15). The prevailing case-control approach to understanding the relationships between neurophysiology and psychopathology is hindered by flaws in traditional categorical approaches to mental health diagnoses, including arbitrary boundaries between pathology and normality, overlap and cooccurrence within diagnostic categories (16). Examples of alternative approaches to the case-control paradigm emphasise data-driven patterns of co-occurrence between dimensional mental health symptoms (17; 18; 19), or neurocognitive test scores (20).

Departing from the case-control approach, we can identify data-driven subgroups of individuals and examine differences between subgroups. By starting with analyses of brain activity to characterise individuals before looking at health outcomes, we have greater opportunity to build robust models and generate novel hypotheses from an empirical perspective. The methodological pipeline developed in this work offers a novel, flexible and scaleable approach for identifying subgroups of individuals on the basis of resting state EEG measurements. This builds on a nascent but growing body of work that uses data-driven analysis of EEG as a starting point for investigating the relationships between neurophysiology, health and functional outcomes (21; 22; 23). To enable development of these data-driven insights and improve our understanding of the relationships between brain characteristics and mental health during adolescent development, there is a need for population-based studies to obtain large datasets on these factors together in this understudied age range.

The Longitudinal Adolescent Brain Study (LABS), being conducted at the Thompson Institute in Queensland, Australia, is a longitudinal cohort study examining the interactions between environmental and psychosocial risk factors, and outcomes including cognition, self-report mental health symptoms, neuroimaging measures, and psychiatric diagnoses (24). The aims for LABS include: identifying early manifestations of psychopathology in young people aged 12 - 17 years; monitoring neurophysiological changes and their relationships with external factors and mental health outcomes over this developmental period; and enabling better identification of risk and early intervention to improve cognitive development and mental health trajectories for young people. The present study uses data collected from the baseline timepoint in LABS. Further information on data collection and study protocols for LABS are provided in the Methods section. In this paper we aim to identify data-driven subgroups of LABS participants using EEG data.

To identify subgroups of adolescents based on EEG characteristics, we implement a selection of popular unsupervised clustering algorithms. Unsupervised clustering methods are a useful set of statistical tools for identifying subgroups within data, without requiring prior information about labels of classifications of groups to which individual data points are expected to belong (25). Different clustering algorithms will emphasise or be sensitive to different aspects of input data in their clustering solutions, so it is valuable to consider results from multiple clustering algorithms especially when there is limited prior evidence about any subgroups that are expected to exist within a dataset. Comparing results across multiple algorithms allows identification of well-separated subgroups with more confidence and enables inference that is less sensitive to choice of method, as opposed to choosing one ‘best’ model.

In the context of the literature introduced here it is evident that to better understand the relationships between brain activity, cognitive function and mental health risk in adolescence, there is value in identifying data-driven subgroups based on brain measurements and comparing cognitive function and mental health outcomes between these subgroups. The present study had several goals: (i) using frequency information from resting state EEG data collected at the baseline timepoint in LABS, to cluster 12-year-olds into subgroups; (ii) to identify stable, well-separated subgroups of individuals across different clustering methods; (iii) to interpret the EEG characteristics differentiating these subgroups; (iv) and to test for differences in cognitive function, mental health symptoms and other health indicators between these EEG-based clusters.

With these goals in mind, we aimed to address the following research questions:

1. Using popular unsupervised clustering algorithms, how can we develop a flexible analysis pipeline to identify subgroups of individuals based on frequency information from resting state EEG?
2. Across clusters of individuals, what characteristic profiles of resting state EEG content differentiate these clusters?
3. How do individuals across clusters vary in terms of psychological distress, wellbeing, sleep quality, and cognitive function?
4. Are these data-driven neurophysiological groups associated with different risk or protective profiles for mental health and cognitive function?

## 3 Methods

### 3.1 The Longitudinal Adolescent Brain Study

#### 3.1.1 Participants

The present study uses cross-sectional data collected from a community sample of young people at baseline entry to the Longitudinal Adolescent Brain Study conducted at the Thompson Institute, University of the Sunshine Coast. Initiated in July 2018, LABS collects participant data every 4 months over 5 years, following participants from ages 12 to 17 — totalling 15 timepoints for each participant. At each timepoint, data collected includes: a self-report questionnaire on demographic information, mental health and wellbeing measures; a neuropsychiatric interview; a battery of cognitive assessments; and neuroimaging scans including resting state and task-based EEG, and structural and functional MRI. Inclusion criteria are being 12 years of age and in Grade 7 (the first year of secondary school) at the time of enrolment to participate. Exclusion criteria include neurological disorder diagnosis, intellectual disability, major illness, or head injury with unconsciousness exceeding 30 minutes.

Further details on data collection and study protocols for LABS are available in previous publications (24; 26).

#### 3.1.2 Ethical Approval

LABS received ethical approval from the University of the Sunshine Coast Human Research Ethics Committee (Approval Number: A181064). Written informed assent and consent was obtained from all participants and their guardian/s. For data analysis conducted at the Queensland University of Technology (QUT), the QUT Human Research Ethics Committee assessed this research as meeting the conditions for exemption from HREC review and approval in accordance with section 5.1.22 of the National Statement on Ethical Conduct in Human Research (2007). Exemption Number: 2021000159.

#### 3.1.3 Data collection

##### EEG Data

For EEG recordings, participants were fitted with a Biosemi headcap mounted with 32 Ag/AgCl active electrodes and arrayed in a 10-10 configuration at a sampling rate of 1024 Hz using a Biosemi ActiveTwo system. To monitor eye movements, horizontal electrooculogram (EOG) was recorded using electrodes placed one-centimetre lateral to the outer canthus of each eye, while vertical EOG was recorded using supra- and infraorbital electrodes on the left eye. Each electrode was filled with a conductive gel and DC offsets were checked to assess electrode contact quality before recording. Eyes-closed, resting state data was recorded for 4 minutes, and participants were instructed to rest and minimise eye movements during the recording. For the present study we analysed data recorded from the vertex electrode (Cz) only, as we were not aiming to investigate patterns across different brain regions.

##### Mental Health & Cognitive Function Measures

In the present study we examine a variety of outcomes related to mental health, collected by self-report questionnaire in LABS. These include: psychological distress as measured by the K10 (27), and SPHERE-12 (28); Suicidal ideation measured by the Suicidal Ideation Attributes Scale (SIDAS) (29); Wellbeing measured by the COMPAS-W (30); and sleep quality, measured by the Pittsburgh Sleep Quality Index (PSQI) (31).

We also examined a number of cognitive function measures from the CogState cognitive test battery including reaction time measures for the Detection (DET), Identification (IDN) and One Back (OBK) tasks, with log10 transformed reaction times for correct responses where lower scores indicate better performance. These measures also included accuracy for the Two Back (TWOB) task using an arcsine transformation of the square root of the proportion of correct responses, for which higher scores indicate better performance (32). This is the standard scoring for these tests. Full details on LABS data collection protocols for these measures are available in previous publications (24).

### 3.2 Statistical analyses

To address the research questions outlined above, this study implemented a multi-stage analysis pipeline. The first stage for clustering based on EEG frequency characteristics included: EEG pre-processing; frequency decomposition with multitaper analysis; and selection and calculation of frequency features. The second stage included: dimensionality reduction using principal components analysis; and applying a number of popular unsupervised clustering algorithms to this dimension-reduced data.

In the third stage to identify well-separated subgroups of participants who are differentiated by characteristic EEG profiles, we selected core clusters of individuals who were consistently grouped together across the different clustering methods. This allowed us to identify well-separated subgroups with distinct profiles of EEG characteristics. After implementing this pipeline and identifying core clusters, we described the principal components and EEG frequency characteristics that differentiate these core clusters. Finally, we applied Bayesian regression models with cluster allocation as a categorical predictive variable to examine between-cluster differences in these health and cognitive function measures. Expected marginal means for pairwise contrasts between clusters were calculated from these regression models to identify the magnitude and associated uncertainty for pairwise differences in these measures between the clusters.

#### 3.2.1 Pre-processing

EEG data tend to be characterised by non-stationarity, a low signal-to-noise ratio, and artifacts from a number of sources including endogenous biological signals, background electrical interference, and variable electrode contact quality (33). In this study we implemented an automated data cleaning and pre-processing pipeline, using a variety of tools available through the EEGLAB toolbox in MATLAB (34), including: ICLabel using a trained algorithm to probabilistically label noise components from Independent Component Analysis (ICA) (35); Adaptive Mixture ICA (AMICA), which is demonstrated to be an effective variant of ICA for EEG artifact removal (36); and Artifact Subspace Reconstruction (ASR) (37). An automated approach was implemented to enable efficient and consistent pre-processing for EEG data across a larga number of participants, and to be consistent with the flexible and scaleable design approach of this analysis pipeline. Further details on the automated EEG cleaning and pre-processing pipeline are presented in the Supplementary Materials.

#### 3.2.2 Multitaper Analysis & selection of frequency features

To study frequency content of EEG data we used multitaper analysis, a frequency decomposition method that is suitable for non-stationary signals and offers good frequency specificity (38). This is a popular technique for frequency analysis that has been used widely in recent EEG research literature (39; 40), and offers an improved signal-to-noise ratio for detecting rhythmic activity in a signal relative to other standard frequency decomposition methods (41; 42).

Multitaper analysis was conducted using the Chronux toolbox in MATLAB (43). This method enabled us to calculate power spectral densities with good frequency resolution and minimal ‘bleeding’ of power across adjacent frequency bands (44). Multitaper time-frequency spectrograms were calculated from resting state, eyes closed EEG recordings, and were then averaged over time to yield power spectral densities (PSDs) representing the average frequency content of EEG activity for each individual LABS participant during the recording session.

From these PSDs we calculated a selection of 8 commonly used summary metrics to capture important characteristics of individual EEG frequency content. Individual Alpha Frequency (IAF) was calculated as the frequency at which maximum power is observed in the range 7-14 Hz (45; 46), which is known to typically increase through adolescence before reaching a peak frequency that has a typical mean of around 10 Hz in adulthood (47). 95% Spectral Edge Frequency (SEF95) was calculated to identify the frequency below which 95% of the power for a PSD is contained, and indicative of the degree of dispersion in the PSD (48). Spectral entropy was calculated as the Shannon entropy of the self-normalised PSD (49), implemented with the ForeCA package in R (50; 51). Finally we included fractional power in bands of width 4 Hz between 0 and 16 Hz, and total power for the PSD. In adults, typical frequency bands and their approximate boundaries are delta (0.5-4 Hz), theta (4-8 Hz), alpha (8–14 Hz), beta (14–30 Hz), and gamma (over 30 Hz) (52). For ease of interpretation we include these traditional frequency band labels as an approximate heuristic guide for labeling the fractional power features we used in 4 Hz intervals up to 16 Hz (low beta), though these bands are not uniformly representative of discrete functional categories of EEG activity in children and adolescents (53).

#### 3.2.3 Principal Components Analysis

From the frequency features calculated from the results of multitaper analysis, we conducted dimensionality reduction with PCA using the prcomp() function from the ‘stats’ package in R (54). We identified dimensionality reduction as an important step in this analysis pipeline, for several reasons. Due to the small number of participants in this dataset, inclusion of this dimensionality reduction step enabled flexibility and scalability of this method to accommodate a large number of input features and avoid some of the challenges around clustering high-dimensional data (55). It is also important to improve model parsimony and interpretability for the clustering stages of this work. The PCA step enabled flexibility of this method to accommodate higher-dimensional input data in future, for example by looking at a large number of summary features across multiple EEG channels to look at activity patterns across multiple brain regions. We chose to retain 3 principal components based on finding an elbow in the scree plot, which coincided with eigenvalues greater than 1. Looking at the principal component coefficients representing the input features in a lower dimensional space, we were able to identify the combinations and contrasts in EEG features that account for the most variance between individuals.

#### 3.2.4 Unsupervised clustering

Based on the principal component scores from the previous stage of analysis, we implemented 3 popular unsupervised clustering algorithms to identify data-driven subgroups of individuals on the basis of EEG frequency characteristics. These algorithms were *k*-means, hierarchical clustering (HC) using Ward’s method, and Gaussian mixture model (GMM), each of which has different assumptions and sensitivities and provides different perspectives on subgroup structure within the data. We calculated results for *k*-means using the kmeans() function with default settings from the base package ‘stats’ in R (54). We calculated results for hierarchical clustering using the hclust() function with method “ward.D2” from the base package ‘stats’ in R (54). We calculated results for GMM with default settings including Euclidean distance and a diagonal covariance matrix using the package ‘ClusterR’ in R, version 1.22 (56). For the applications in this work, we use the crisp projection of GMM results with allocation of each data point to the mixture component in which it is most likely to belong. Further details on each algorithm are presented in the Supplementary Materials.

For each clustering algorithm, the optimal number of clusters *k* was selected on the basis of a number of internal validation indices which could be calculated for all three methods. Internal validation indices can be used to identify the number of clusters that creates the most compact and well-separated set of subgroups in the data (25). Each index is calculated based on slightly different metrics, so using a selection of multiple indices can be more robust than relying on a single index. Selected indices included the Dunn index (57), silhouette coefficient (58), Davies-Bouldin index (59), Calinski-Harabasz index (60), and Xie-Beni index (61). Clustering internal validation indices were calculated in R using the clusterCrit package, v1.2.8 (62).

#### 3.2.5 Identifying core clusters across methods

To identify subgroups of individuals that were consistent across the clustering methods implemented, we selected ‘core’ clusters that co-occurred in all three sets of results. As the labels and locations of clusters were not consistent across methods, we identified overlapping membership in the unique combinations of cluster labels across the 3 methods. We retained core clusters with the highest overlapping membership across all methods, aiming for a set of combined results where each cluster label from each method was represented only once in the final set of core clusters.

This process was undertaken to identify well-separated subgroups of individuals and enable inferences about members of these core clusters that are more robust and less sensitive to the choice of clustering method. This also enabled identification of clearly differentiated subgroups with distinct profiles of EEG characteristics. There are some limitations to this simple approach for integrating multiple sets of clustering results, in that individuals who are not consistently clustered across different algorithms (representing borderline or edge cases located between the final core clusters) are not allocated to the final set of core clusters. For this application we decided this was a reasonable method, as internal validation criteria supported clustering solutions with a similar structure across the three algorithms and only a small number of participants (n = 11) were not allocated to the core clusters. This enabled us to identify final subgroups with clearly differentiated profiles of EEG characteristics. Further details on the intersection of clustering results and selection of core groups are provided in the Supplementary Materials.

#### 3.2.6 Testing for differences in cognitive function & health measures between core clusters

To examine patterns of difference in levels of psychological distress, wellbeing, sleep quality and cognitive function across the final EEG-based clusters, we implemented Bayesian regression models using the “brms” package in R, version 2.15.0 (63). These models were designed to predict scores on each measure using core cluster allocation as a categorical predictive variable, with the largest cluster (cluster 1) as the reference category and including sex as a covariate. Prior sensitivity analyses were conducted for all Bayesian models, comparing the default improper flat prior on regression coefficients used in “brms” with two alternatives. The alternative priors tested were a normal distribution with mean 0 and standard deviation 10, and a Student’s T distribution with 2 degrees of freedom, location 0 and scale 10. These priors each provided appropriate support, covering the anticipated range of the parameter values for the regression coefficients.

From these Bayesian regression models we calculated posterior probabilities for each outcome measure being greater in one cluster than another, for all pairwise combinations of the clusters. We also calculated expected marginal means for pairwise contrasts between clusters using the “emmeans” package in R, version 1.5.5-1 (64). This allowed us to identify posterior credible intervals for the magnitude of the differences in these health measures between all pairwise combinations of the clusters.

## 4 Results

From cross-sectional baseline data for 12-year-old (M = 12.64, SD = 0.32) participants that had been recruited into LABS when this article was written, data from 59 individuals were included after removing individuals with missing or corrupted EEG data.

### 4.1 Principal Components Analysis

Based on visual inspection of the scree plot and identifying components with an eigenvalue greater than 1, the first 3 principal components were retained which together explain 80.6% of the total variance. Initial visual inspection of scatter plots of individuals by principal component scores indicated that individuals were well dispersed across these 3 components, and some areas of higher and lower density were apparent — suggesting the opportunity for unsupervised clustering methods to identify useful subgrouping structure in this dataset. Further details including eigenvalues, proportions of variance explained and a scree plot are available in the Supplementary Materials.

#### 4.1.1 PCA Interpretation

Table 1 presents coefficients for each of the 8 input frequency features on the 3 retained components. Choosing a heuristic threshold magnitude of 0.4 for these coefficients, we can identify the most influential variables contributing to the variance explained in each component.

**Table 1:** Component coefficients for PCA of frequency features. Coefficients with magnitude *>* 0.4 are in bold. IAF = individual alpha frequency; SEF95 = 95% Spectral Edge Frequency. 4 Hz power bands are as a proportion of total power.

|  | PC1 | PC2 | PC3 |
| --- | --- | --- | --- |
| IAF | -0.2339 | **0.5657** | -0.0179 |
| SEF95 | **0.4128** | 0.0927 | 0.1925 |
| Total Power | -0.3896 | -0.1587 | -0.1604 |
| Delta (0-4 Hz) | **0.5031** | -0.0360 | -0.1218 |
| Theta (4-8 Hz) | -0.0973 | **-0.6179** | 0.2544 |
| Alpha (8-12 Hz) | **-0.4789** | 0.2204 | -0.2257 |
| Low Beta (12-16 Hz) | 0.0885 | **0.4334** | **0.6317** |
| Spectral Entropy | 0.3512 | 0.1635 | **-0.6383** |

Component 1 explains 41% of the overall variance, and represents an average of 95% spectral edge frequency (SEF95) and fractional power in the delta range (0-4 Hz), contrasted with fractional power in the alpha range (8 - 12 Hz). Accordingly individuals with higher scores on this component would typically have a higher SEF95, a higher proportion of power at low frequencies between 0 and 4 Hz, contrasted with a lower proportion power between 8 and 12 Hz. This component can be thought of as describing the level of dispersion and delta power, contrasted with alpha power.

Component 2 explains 27% of the overall variance, and represents an average of Individual Alpha Frequency (IAF) and fractional power in the low beta range (12-16 Hz), contrasted with relative power in the theta range (4-8 Hz). Higher scores on this component identify individuals with higher frequency alpha peaks, higher relative beta power and lower relative theta power. Given the variability of IAF between 7-14 Hz, overlapping with the theta (4-8 Hz) and lower beta (12-16 Hz) bandwidths, this component therefore appears to mainly capture variability related to the frequency location of the alpha peak for each individual.

Component 3 explains 13% of the overall variance, and represents a contrast of relative power in the low beta range (12-16 Hz) with spectral entropy. This component captures the disparity between these variables — indicating that individuals in our dataset with higher relative power between 12-16Hz will tend to have a less complex PSD with lower spectral entropy, and vice versa.

### 4.2 Unsupervised clustering methods

Results from each clustering algorithm are presented below, before identifying integrated core clusters of individuals that are consistently clustered together across clustering methods. These core clusters are then described in terms of their average PSD, principal component scores, and original frequency feature values.

On the basis of the internal validation criteria introduced above, alongside the goals of model parsimony and interpretability for the final integrated clusters, we chose to implement a 5-cluster solution in each of the three individual clustering methods. Examining results from multiple internal validation indices, we identified support for a 5-cluster solution as the optimal choice for the *k*-means and GMM algorithms. For hierarchical clustering, we found marginal support for a 5-cluster or 7-cluster solution being optimal. For the sake of parsimony and interpretability in the final set of core clusters, we decided to implement a 5-cluster solution for all three algorithms to have a cohesive approach across the methods. This allowed us to identify the most consistent structure across results from the different algorithms, and maximise the number of individuals allocated to the integrated core clusters. Further details on selecting the number of clusters *K* for each method are provided in the Supplementary Materials.

Table 2 presents the number of individuals assigned to each cluster label for the three clustering algorithms, and for overlapping ‘core’ clusters. For each algorithm, cluster labels (1-5) have been assigned by decreasing cluster size except for HC clusters 3 and 4, for which labels were switched for the sake of clearer visual comparison across plots.

**Table 2:** Cluster membership for different clustering algorithms & core clusters, *K*=5

| Cluster label | 1 | 2 | 3 | 4 | 5 |
| --- | --- | --- | --- | --- | --- |
| *k*-means | 18 | 15 | 13 | 10 | 3 |
| HC | 17 | 15 | 9 | 15 | 3 |
| GMM | 21 | 13 | 13 | 9 | 3 |
| Core clusters | 15 | 12 | 9 | 9 | 3 |

Figure 2 presents 2D scatter plots of clustering results for each clustering method, displayed according to each pairwise combination of the 3 retained principal components. 3D scatter plots are available in the Supplementary Materials.

**Fig 1.**
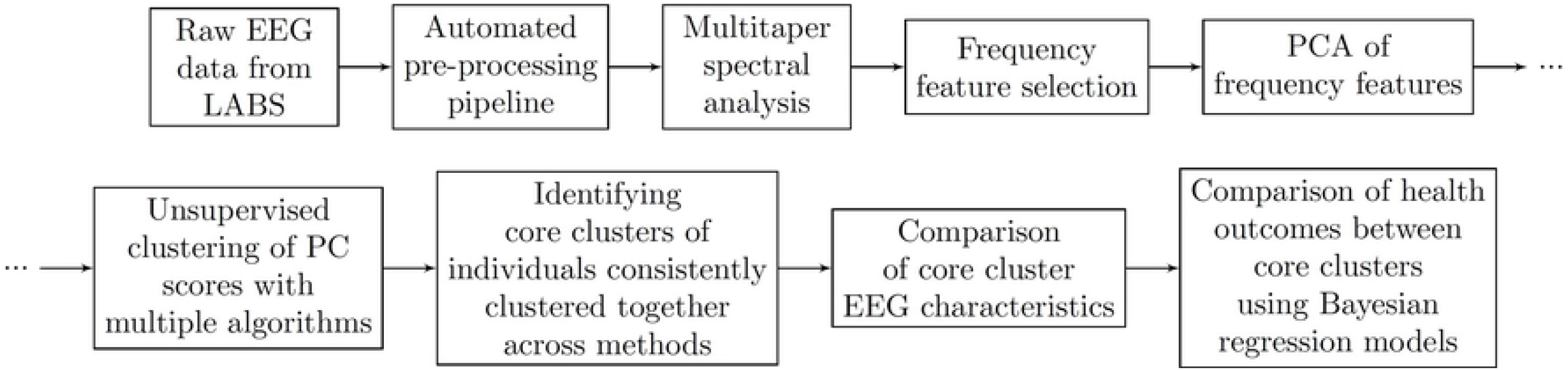
Methods overview block diagram.

**Fig 2.**
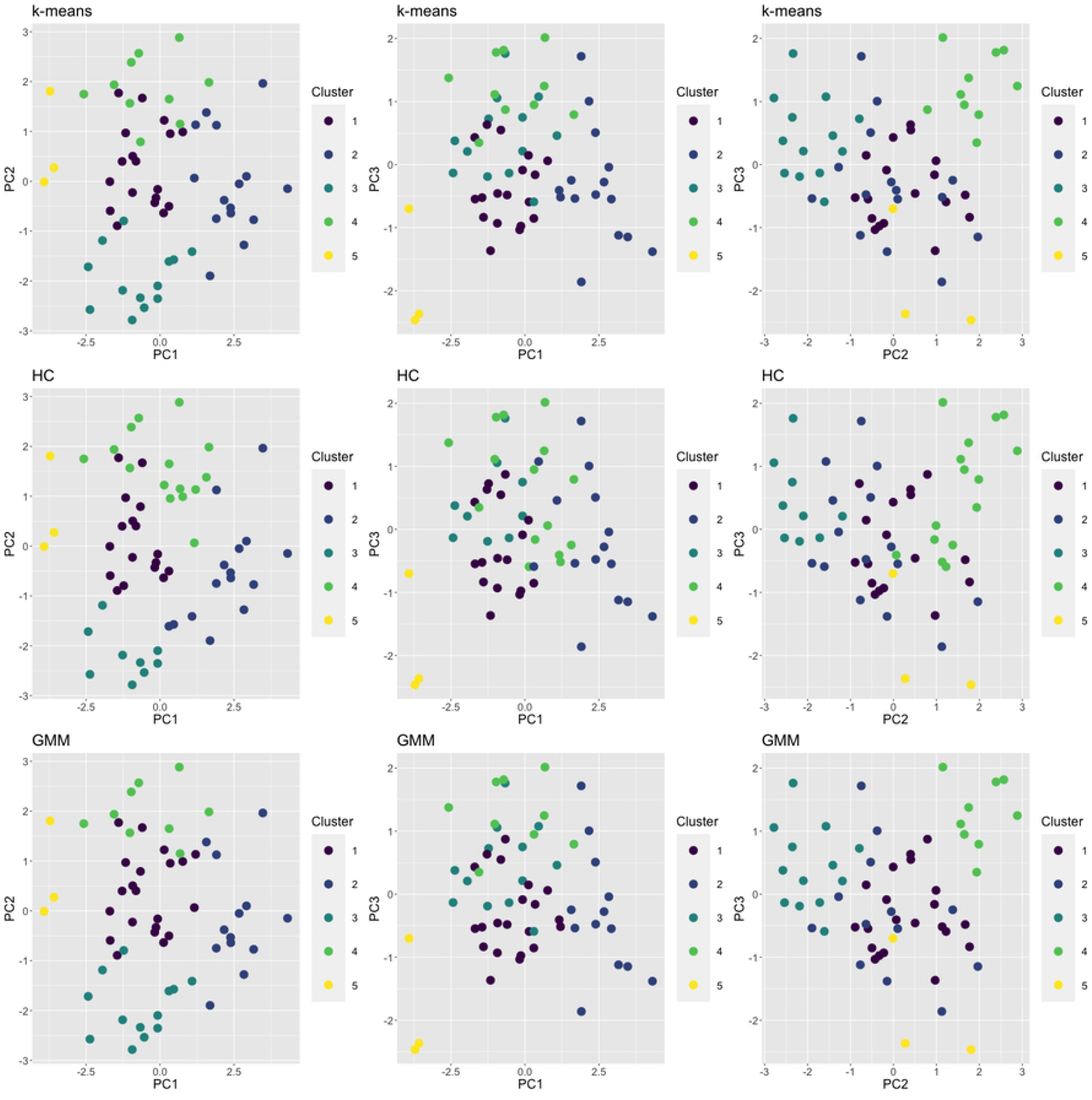
2D scatter plots of individuals by component scores, coloured by cluster membership, *K* = 5. Top row = *k*-means; middle row = HC; bottom row = GMM. Left column = PC1 v PC2; middle column = PC1 v PC3; right column = PC2 v PC3.

### 4.3 Core clusters across methods

As discussed above, we aimed to retain the number of core clusters equal to the minimum *K* chosen across clustering methods, where each core cluster represents an intersection across a unique combination of cluster labels from each method, and cluster labels from each method were represented only once in the core clusters. In total we retained 5 core clusters which included a total of 48 individuals, and 11 individuals were not allocated.

Figure 3 presents 2D scatter plots of individuals by principal component scores coloured by core cluster membership after identification of overlapping clustering results between the algorithms, displayed by each combination of the 3 retained principal components. Individuals who were not included in any core cluster as they were not clustered consistently between methods (*n* = 11) have been removed from these plots.

**Fig 3.**
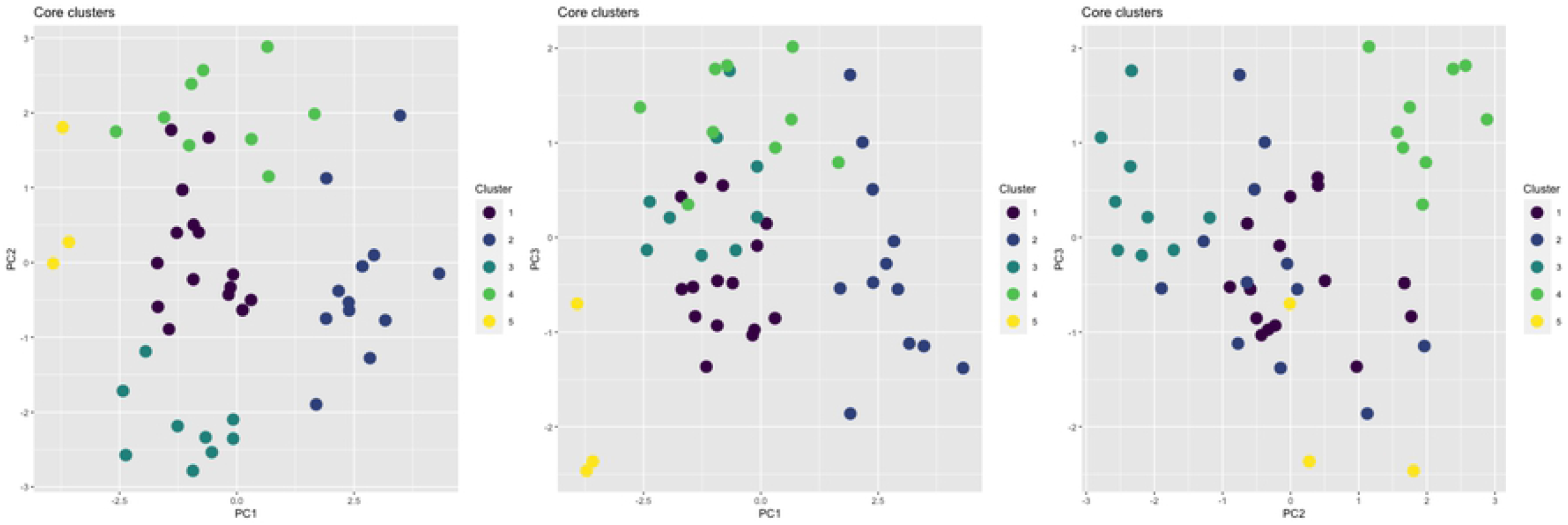
2D scatter plots of individuals by component scores, coloured by core cluster membership, *K* = 5. Individuals not allocated to core clusters have been removed. Left panel = PC1 v PC2; middle panel = PC1 v PC3; right panel = PC2 v PC3.

#### 4.3.1 Core clusters - principal component scores

Looking at the cluster locations in Figure 3 and the mean principal component scores for each core cluster in Table 3, we can identify the frequency characteristics that are expected to differentiate the core clusters.

**Table 3:** Core cluster profiles: Principal component scores. Mean (SD)

| Core cluster | PC1 | PC2 | PC3 |
| --- | --- | --- | --- |
| 1 | -0.7938 (0.6658) | 0.1293 (0.8179) | -0.4220 (0.6231) |
| 2 | 2.6522 (0.7516) | -0.2716 (1.0223) | -0.3466 (1.0275) |
| 3 | -1.1448 (0.9158) | -2.1971 (0.4879) | 0.4344 (0.6506) |
| 4 | -0.3940 (1.3165) | 1.9860 (0.5424) | 1.2704 (0.5398) |
| 5 | -3.7381 (0.1681) | 0.6892 (0.9771) | -1.8443 (0.9915) |

Cluster 1 was the most central, with small scores on every PC, so it was not obvious from the PC scores how frequency characteristics for this cluster were expected to differ from the others. Clusters 2, 3 and 5 had high magnitude scores on PC1, and were expected to differ by their level of frequency dispersion and low-frequency power, contrasted with the magnitude of the individual alpha peak. Clusters 3 and 4 had high magnitude scores on PC2, differentiated by the frequency location of the individual alpha peak, and by higher frequency activity in the range 12-16Hz. Clusters 4 and 5 had high magnitude scores on PC3, differing by the level of contrast between relative power in the range 12-16 Hz and spectral entropy.

We can further investigate these patterns in cluster principal component scores by looking at mean scores on the original frequency features (Table 4), and mean power spectral density plots (Figure 4) for each core cluster.

**Table 4:**
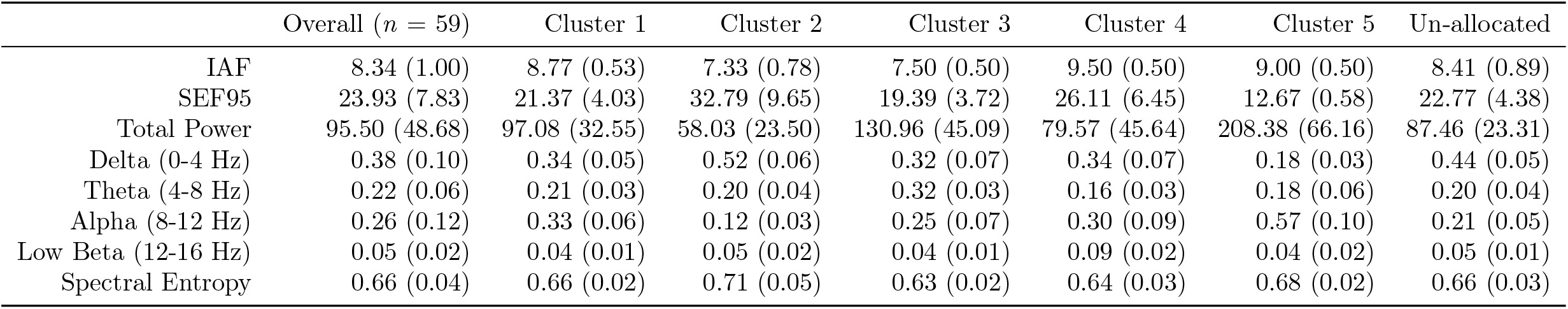
Descriptive statistics for EEG features. The first column presents statistics for all participants (*n* = 59), and the remaining columns present statistics for individuals allocated to each of the final core clusters. IAF = individual alpha frequency; SEF95 = 95% Spectral Edge Frequency. 4 Hz power bands are as a proportion of total power.

**Fig 4.**
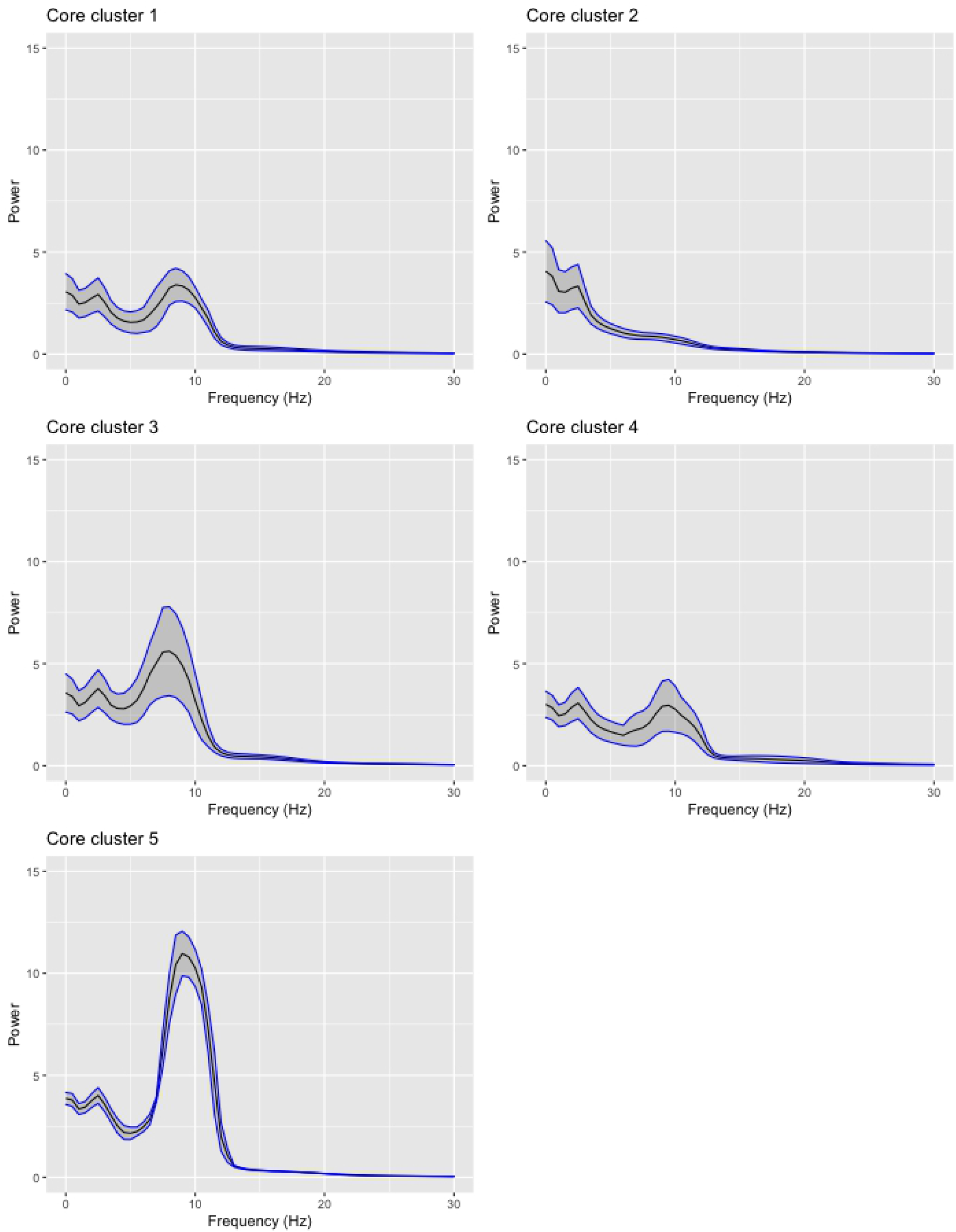
Mean PSDs for each core cluster. Shaded regions represent standard deviations.

#### 4.3.2 Core clusters - EEG frequency features

From Table 4 and Figure 4, and as expected from the principal component scores, cluster 1 (n=15) sits close to the middle of the range for each of the frequency features calculated. It has a well defined alpha peak and a moderate IAF (8.77) relative to other clusters.

We can see that cluster 2 (n=12) is characterised by noticeably lower theta and alpha activity in the range 4-12 Hz, high power at low frequencies, high dispersion, and low total power. Members of this cluster have high relative power at low frequencies compared to other clusters, with the lowest IAF (7.33) and more than half of their relative power between 0-4 Hz (0.52), contrasted with having the lowest relative alpha power between 8-12 Hz (0.12). Cluster 2 is also characterised by high dispersion with the highest SEF95 (32.79) and spectral entropy (0.71) indicating a complex frequency distribution with power distributed across a wide range. This cluster also has the lowest total power (58.03).

Cluster 3 (n=9) has high power at low frequencies, with low dispersion and a pronounced alpha peak. With a low IAF (7.50) and the highest relative power in the range 4-8 Hz (0.32), we can identify high theta activity, and a pronounced alpha peak at the low range for IAF. A low SEF95 (19.39) further indicates that most of the power is concentrated at low frequencies for this cluster. Cluster 3 also has low spectral entropy (0.63) and the equal lowest relative power between 12-16 Hz (0.04), indicating a less complex frequency distribution that is concentrated at lower frequencies.

Cluster 4 (n = 9) has a high frequency alpha peak with the highest IAF (9.50) and the lowest relative power in the range 4-8 Hz (0.16), indicating low theta activity. It also distinguished by having the highest relative power in the range 12-16 Hz (0.09), demonstrating higher activity than other clusters in the upper alpha - lower beta range.

Cluster 5, the smallest cluster (n = 3), has high scores on principal component 1 and is characterised by a high magnitude, high frequency alpha peak, low dispersion, and high total power. It has a high IAF (9.00) and the highest activity in the alpha range 8-12 Hz (0.57), with over half its relative power occurring in this range. This cluster also has the highest total power with more than double the mean total power across all participants (208.38), the lowest relative delta power of any cluster in the range 0-4 Hz (0.18) and the lowest SEF95 (12.67), indicating a distinct and high degree of power concentrated in the alpha range.

### 4.4 Comparing cognitive function & mental health measures across clusters

Descriptive statistics for EEG frequency features in each cluster are presented in Table 4. Descriptive statistics for mental health and cognitive function measures in each core cluster are presented in Table 5. Figure 5 presents column graphs showing Z-scores based on the global mean and standard deviation for each EEG feature (top panel) and each health and behavioural measure (bottom panel) across the 5 clusters, allowing a comparison of the EEG and health characteristics in each group. Violin plots showing the distribution of scores for health measures in each cluster are available in Supplementary Figure 5.

**Table 5:** Descriptive statistics for mental health and cognitive function measures. The first column presents statistics for all participants (*n* = 59), and the remaining columns present statistics for individuals allocated to each of the final core clusters. K10 = Kessler Psychological Distress Scale; SPHERE12 = SPHERE-12 Screening Questionnaire; SIDAS = Suicidal Ideation Attributes Scale; COMPAS-W = COMPAS-W Wellbeing Scale; PSQI = Pittsburgh Sleep Quality Index; DET – RT = CogState Detection Task Reaction Time; IDN - RT = CogState Identification Task Reaction Time; OBK - RT = CogState One Back Task Reaction Time; TWOB – Acc = CogState Two Back Task Accuracy Score.

| Measure | Overall | Cluster 1 | Cluster 2 | Cluster 3 | Cluster 4 | Cluster 5 |
| --- | --- | --- | --- | --- | --- | --- |
| N | 59 | 15 | 12 | 9 | 9 | 3 |
| Age (weeks), mean (sd) | 657.11 (16.48) | 655.71 (15.49) | 659.19 (17.26) | 658.81 (16.52) | 659.05 (19.12) | 653.14 (23.94) |
| Female, $n$ (%) | 28 (47.46) | 8 (53.33) | 4 (33.33) | 6 (66.66) | 3 (33.33) | 1 (33.33) |
| K10, mean (sd) | 15.66 (4.63) | 14.40 (4.29) | 16.50 (4.40) | 14.33 (2.83) | 14.78 (5.54) | 16.67 (2.31) |
| SPHERE12, mean (sd) | 4.02 (3.63) | 2.80 (3.10) | 5.42 (3.26) | 4.67 (3.32) | 3.33 (4.00) | 2.00 (1.73) |
| SIDAS, mean (sd) | 3.17 (8.44) | 2.47 (6.36) | 1.42 (4.01) | 8.89 (17.00) | 2.00 (4.69) | 0.00 (0.00) |
| COMPAS-W, mean (sd) | 102.17 (10.95) | 105.00 (10.11) | 99.67 (11.98) | 99.89 (12.94) | 105.89 (10.35) | 108.33 (6.43) |
| PSQI, mean (sd) | 4.95 (2.81) | 4.80 (2.27) | 4.75 (1.66) | 6.67 (2.92) | 3.75 (2.43) | 2.67 (1.15) |
| DET - RT, mean (sd) | 2.54 (0.09) | 2.57 (0.08) | 2.50 (0.03) | 2.58 (0.19) | 2.50 (0.03) | 2.58 (0.01) |
| IDN - RT, mean (sd) | 2.73 (0.07) | 2.76 (0.08) | 2.69 (0.04) | 2.78 (0.08) | 2.70 (0.05) | 2.71 (0.03) |
| OBK - RT, mean (sd) | 2.89 (0.09) | 2.92 (0.09) | 2.87 (0.11) | 2.94 (0.11) | 2.85 (0.07) | 2.83 (0.05) |
| TWOB - Acc, mean (sd) | 1.25 (0.19) | 1.29 (0.18) | 1.21 (0.14) | 1.09 (0.27) | 1.36 (0.15) | 1.20 (0.12) |

**Fig 5.**
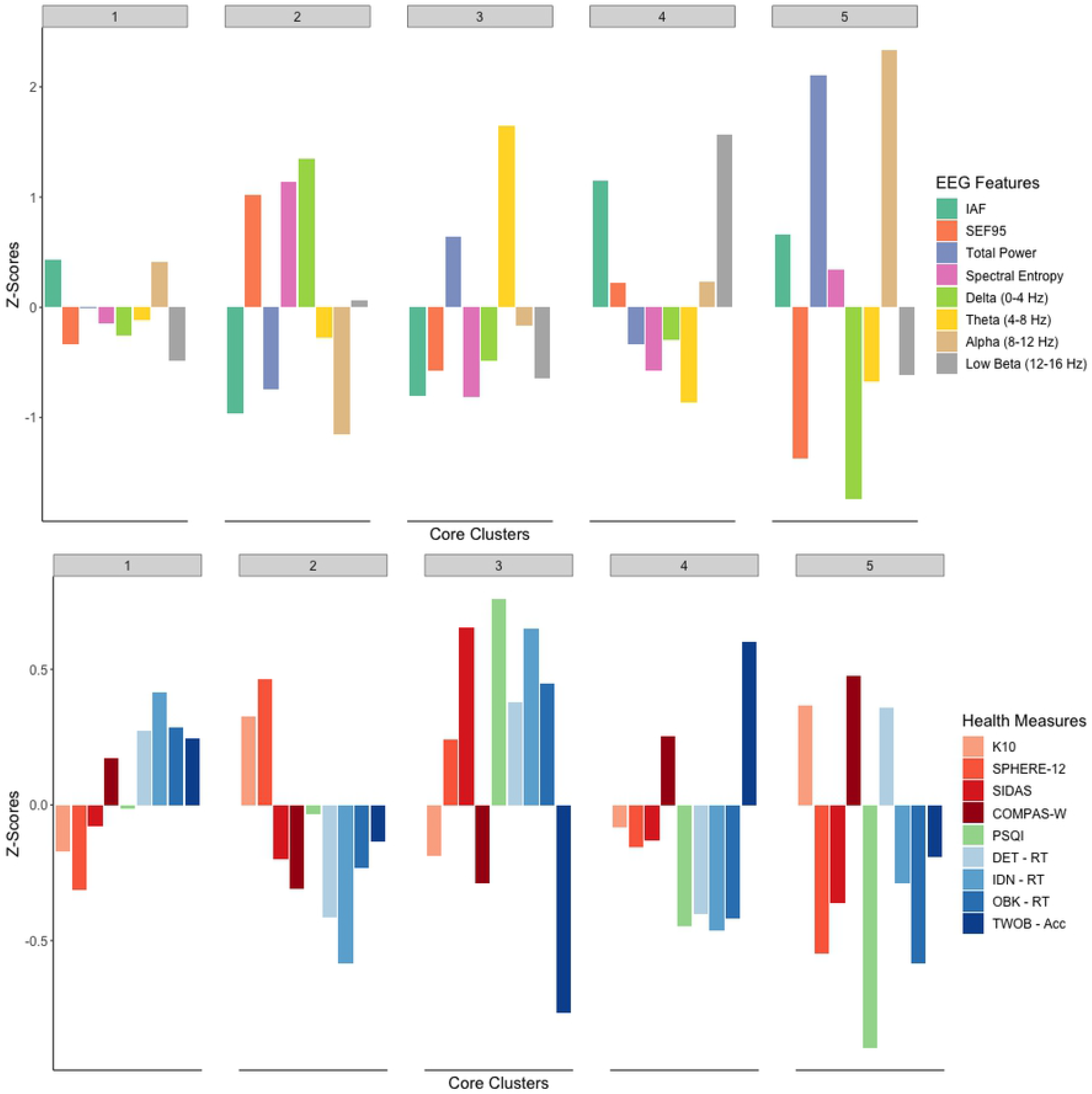
Bar plots showing Z-scores for each EEG feature (top panel) and each health and cognitive measure (bottom panel) across core clusters. IAF = individual alpha frequency; SEF95 = 95% Spectral Edge Frequency. 4 Hz power bands are as a proportion of total power. K10 = Kessler Psychological Distress Scale; SPHERE12 = SPHERE-12 Screening Questionnaire; SIDAS = Suicidal Ideation Attributes Scale; COMPAS-W = COMPAS-W Wellbeing Scale; PSQI = Pittsburgh Sleep Quality Index; DET – RT = CogState Detection Task Reaction Time; IDN - RT = CogState Identification Task Reaction Time; OBK - RT = CogState One Back Task Reaction Time; TWOB – Acc = CogState Two Back Task Accuracy Score.

Bayesian regression models were applied to investigate detailed patterns of difference in cognitive function and health measures between clusters. From sensitivity analyses, the results were not substantially different according to the prior used. Figure 6 presents posterior probabilities for these outcome measures being different between each pairwise combination of clusters. These posterior probabilities are approximated from the models results by calculating the proportion of Markov Chain Monte Carlo iterations in which the condition is true. In Figure 6, red cells indicate greater than 90 % probability that the corresponding measure was higher in cluster A than cluster B (A *>* B). Blue cells indicate greater than 90 % probability that the measure was lower in cluster A than cluster B (A *<* B). Expected marginal means from these models for contrasts between clusters are presented in the Supplementary Materials (Supplementary Table 2), providing information about the magnitude of the differences highlighted in Figure 6.

**Fig 6.**
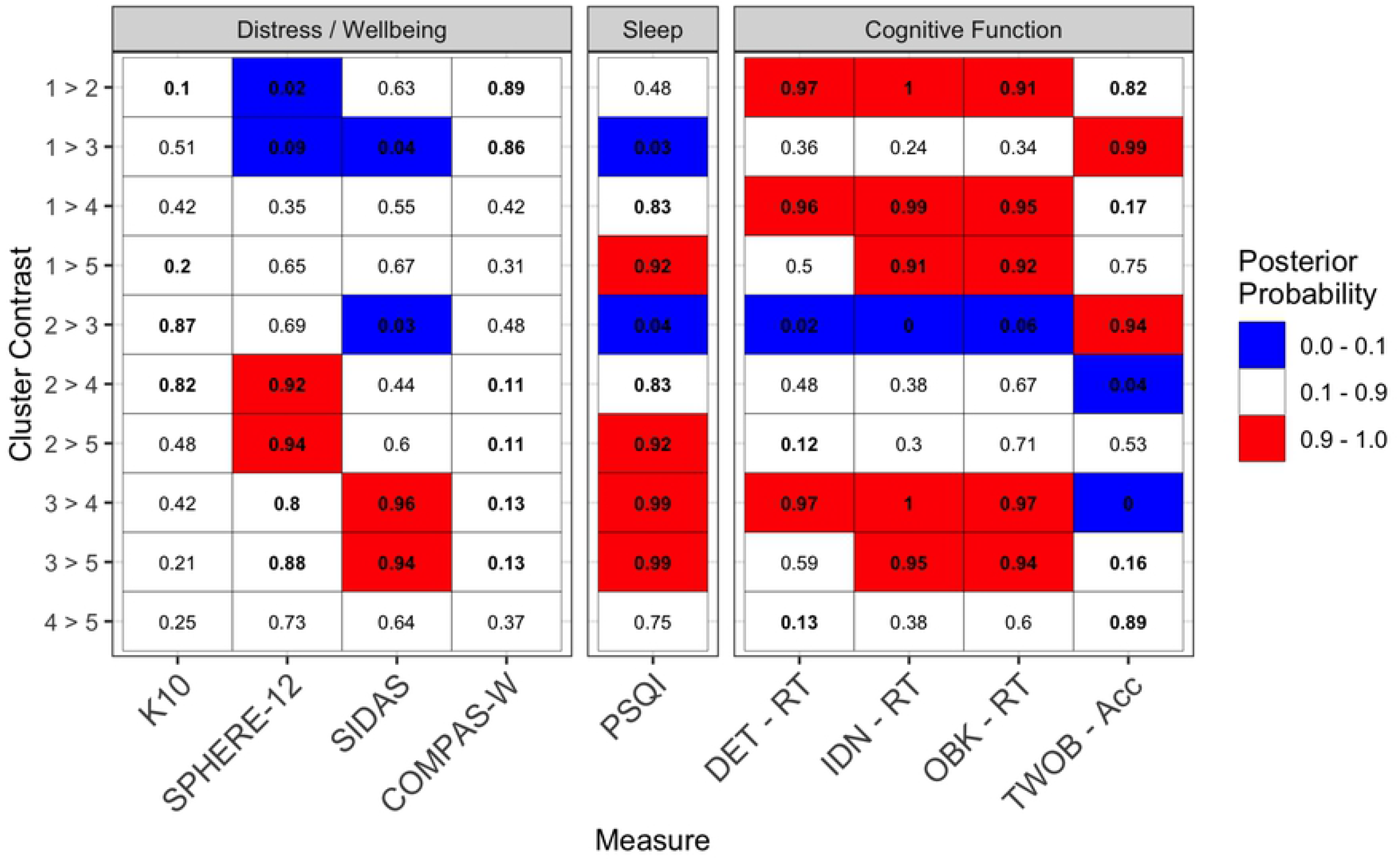
Posterior probabilities for differences in health measures between clusters, calculated from Bayesian regression models. Red cells = greater than 90% probability that measure is higher in cluster A than cluster B (A *>* B). Blue cells = greater than 90% probability that measure is higher in cluster B than cluster A (B *>* A). Bold typeface identifies cells with greater than 80% or less than 20% probability. K10 = Kessler Psychological Distress Scale; SPHERE12 = SPHERE-12 Screening Questionnaire; SIDAS = Suicidal Ideation Attributes Scale; COMPAS-W = COMPAS-W Wellbeing Scale; PSQI = Pittsburgh Sleep Quality Index; DET – RT = CogState Detection Task Reaction Time; IDN - RT = CogState Identification Task Reaction Time; OBK - RT = CogState One Back Task Reaction Time; TWOB – Acc = CogState Two Back Task Accuracy Score.

For suicidality, cluster 3 appeared to have a higher score on the SIDAS scale compared to every other cluster. However, this signal disappeared when one outlying subject in cluster 3 was removed from the analysis, who had the maximum possible SIDAS score, and was also an outlier on reaction time for the CogState detection task (DET - RT). Therefore, findings on between-cluster differences for suicidality (SIDAS) and reaction time on the detection task (DET - RT) should be interpreted with caution, as they are sensitive to the contribution of this one individual. Given that this is a population sample and instances of extreme responding and individuals lying in the tails of distributions are expected occurrences, we decided to include this individual for analyses in the body of this paper.

#### 4.4.1 Cluster Profiles

Looking at the differences in EEG features, health and cognitive function measures across clusters, we could identify tentative EEG and health profiles that may be meaningful in the context of risk prediction for mental health and cognitive development based on characteristic resting state EEG profiles in adolescents. These EEG-based clusters were substantively different in terms of their self-reported psychological distress, wellbeing, sleep quality, and several cognitive function measures. Figures 5-6 and Tables 4-5 show how these groups exhibit meaningful differences in their resting state EEG profiles alongside health measures of interest.

Cluster 1 was the largest cluster, and sat close to the global mean for this sample on most EEG features (Figure 5). From our Bayesian regression models (Figure 6), we identified a high (*>* 90%) probability that members of cluster 1 had lower distress than cluster 2 (K10 and SPHERE-12), lower distress and suicidality than cluster 3 (SPHERE-12 and SIDAS), and slower reactions on all three reaction time tasks than clusters 2 and 4.

Cluster 2 was characterised by low relative theta and alpha power (4-12 Hz), a low frequency alpha peak, and low total power. We found a high probability that members of cluster 2 were more distressed than clusters 1, 4 and 5 (SPHERE-12), and had faster reaction times on all three reaction time tasks than clusters 1 and 3.

Cluster 3 was characterised by a low frequency alpha peak, high relative theta power, and a concentration of power at lower frequencies. Members of cluster 3 had the most concerning sleep and cognitive function profile, with substantially poorer sleep quality than members of all other clusters, slower reaction times than clusters 2, 4 and 5 on reaction time measures, and poorer accuracy on the Two-Back task than clusters 1, 2 and 4. This cluster appeared to have higher suicidality (SIDAS) than other clusters, though as discussed above this signal disappeared when removing one outlying participant, so should be interpreted here with caution. The signals for differences in psychological distress and wellbeing measures were not as strong, but there was moderate evidence (*>*80% probability) to suggest elevated distress (SPHERE-12) and lower self-reported wellbeing (COMPAS-W) in this group relative to clusters 1, 4 and 5. This could suggest that the characteristic resting state EEG profile of cluster 3 is associated with elevated risk of poor outcomes in the domains of mental health, sleep quality and cognitive function in early adolescence.

Cluster 4 was characterised by a high frequency alpha peak, low relative theta power (4-8 Hz), and high relative power in the upper alpha/lower beta range (12-16 Hz). We found moderate to high probability that cluster 4 had lower distress (SPHERE-12) and higher wellbeing (COMPAS-W) than individuals in clusters 2 and 3. Cluster 4 also had moderate (*>* 80%) probability of reporting better sleep quality than clusters 1, 2 and 3. Members of cluster 4 had faster reaction times than clusters 1 and 3 on all three reaction time measures, as well as having higher accuracy than clusters 2 and 3 on the Two-Back task. This could suggest that the EEG characteristics of cluster 4 are associated with improved outcomes in the domains of mental health, sleep quality and cognitive function.

Cluster 5, having only 3 members, shows more variability across these measures. It is characterised by a uniquely high magnitude alpha peak, low dispersion, and high total power. Particular caution should be taken for interpreting results about differences in health and cognitive function for this group, given the very small number of individuals represented. Our models identified high (*>*90%) probability that this cluster had better sleep quality than clusters 1, 2 and 3, and faster reaction times than clusters 1 and 3 on two of the reaction time tasks.

It is also important to note that the relationships between EEG characteristics and health measures were not directly modeled in a regression framework here, and instead the patterns of difference in health measures here are related to the particular combinations of EEG characteristics that distinguish these clusters. The relationships found between individual EEG features and health measures might be quite different if modelling them together as predictor and response variables respectively, but that was not the goal of the present analysis.

These are interesting initial findings that warrant further research and demonstrate the utility of the data-driven method implemented here. Further analyses, with greater statistical power once more data becomes available, will allow us to determine whether there is a robust association between these EEG characteristics and patterns of risk for mental health and cognitive function, cross-sectionally and over time.

## 5 Discussion

In this work we have addressed several main objectives: to develop and validate a flexible analysis pipeline for identifying data-driven subgroups of adolescents based on resting state EEG data; interrogating the patterns of EEG characteristics that distinguish these clusters; to investigate patterns of difference in mental health and cognitive function measures between clusters; and to identify preliminary risk and protective profiles of health and functional measures associated with EEG-based clusters.

With the analysis pipeline developed here, using data-driven unsupervised clustering we were able to identify well-separated subgroups of 12-year-olds on the basis of frequency content from resting state, eyes closed EEG data. While these results are based on measurements from a single EEG electrode, this pipeline has been developed with flexibility and scaleability in mind, and can easily accommodate a larger and/or different set of EEG features calculated across multiple electrodes. The implementation of a Bayesian approach for comparison of health measures between clusters enabled us to make probabilistic statements about between-group differences with quantified uncertainty. This has attractive benefits for application in a clinical setting, where intuitive probabilistic interpretation of risk is desirable for credibility and interpretation of the models for clinicians, and communication of probabilities to healthcare consumers.

The dimensionality reduction stage we have implemented here with PCA enables this analysis framework to accommodate working with data from a small number of individuals, and a large number of features calculated from EEG data. In future applications this dimensionality reduction stage will also allow us to look at EEG characteristics across a number of electrodes, examining spatial variation in activity across different brain regions while avoiding some of the challenges arising when analysing high dimensional data. Working in a dimension-reduced space also improves model parsimony and interpretability for unsupervised clustering results.

Particularly influential EEG variables that we identified as distinguishing individuals included: the level of dispersion of power across frequencies, calculated with the 95% spectral edge frequency; relative power in 4 Hz width frequency bands (delta, theta, alpha and beta) up to 16 Hz; and the location of the individual’s alpha peak frequency. We do not claim that in every case an EEG-based cluster will be associated with pronounced differences in mental health or cognitive function, as there is scope for biology and neurophysiology to vary between individuals without necessarily being associated with substantial health or behavioural differences. However, we have found some patterns of association between characteristic profiles of resting state EEG frequency content and health outcomes which warrant further investigation. For instance the indications of a potential at-risk profile associated with the EEG characteristics of cluster 3, or a protective profile associated with cluster 4, support the utility of these methods and suggest avenues for future investigation. It is also interesting that these two clusters which have contrasting and distinct health and cognitive function profiles also exhibit contrasts on several key EEG features. For example compared to other clusters, cluster 3 exhibits a low frequency alpha peak (IAF), high relative theta (4-12 Hz) power, and low relative power in the upper alpha - lower beta range (12-16 Hz), while cluster 4 has a high IAF, low relative theta power, and high relative beta power. These observations are only descriptive at this stage, but future work will allow us to further investigate whether these combined patterns of EEG frequency features could be useful as indicative biomarkers for risk for mental health and cognitive function.

Given the small sample size in the present study, it is important not to over-interpret these results or to assume that they are able to be broadly generalised. Overall, the strength of the signals for between-cluster differences was weaker for self-report distress and wellbeing measures compared to cognitive testing tasks — which is to be expected, given the known ability of cognitive tests to have better inter-subject reliability and power to detect granular differences between individuals than self-report scales (65; 66; 67). However, the large number of substantial signals identified here relating to differences in mental health and cognitive function between EEG-based clusters supports the utility of the methodological pipeline we have implemented.

The present findings demonstrate the potential for this approach to offer new and important insights around EEG profiles and biomarkers that may be useful in a clinical context. This modelling framework enables insight into detailed patterns of difference between distinctive profiles of resting state EEG activity, and associations with sleep quality and cognitive function measures.

We have foregrounded empirical assessment of brain data, creating opportunities for novel insights into the relationships between neurophysiology, mental health and cognitive function. Our approach offers findings that would not be accessible from a traditional case-control approach of comparing neurophysiology between groups of individuals based on clinical diagnostic categories. Case-control approaches to researching the patterns between brain activity and mental health are especially limited for research in adolescence, when patterns of dimensional symptoms are associated with pluripotent risk for different mental disorder outcomes and often overlap across traditional diagnostic categories (2). Considering the pluripotential nature of mental health problems as they are understood to develop and emerge during adolescence, it was beneficial to use the empirical data-driven approach that we implemented here, studying psychopathology measures on a continuum rather than between diagnostic categories. Other literature has demonstrated that this approach offers reduced constraints from separating and contrasting individuals according to clinical diagnostic labels. Stepping beyond the case-control paradigm enables novel understanding of how different characteristics of brain measurements and mental health outcomes relate to each other. This is advantageous for novel hypothesis generation and improving upon known issues with traditional diagnostic categories including comorbidity and difficulties with differential diagnosis, especially during adolescence (14; 16).

The present study contributes to a growing body of research has demonstrated important interconnections between cognitive function, brain characteristics, and mental health throughout adolescent development. There is some evidence that greater cognitive ability in childhood is associated with decreased risk of internalising mental health symptoms through adolescence, and can attenuate the impact of external stressors on risk of mental health problems (68). Higher childhood cognitive ability is also associated with fewer symptoms of anxiety and depression in adult females and increases risk for potential alcohol abuse in adulthood (69). Some recent findings from longitudinal data show some evidence of a causal relationship between depressive symptoms in adolescence predicting poorer cognitive ability in adulthood. This body of literature suggests important interrelationships between cognitive function, psychopathology and brain health in adolescence (70; 71). Our work highlights that to develop a stronger understanding of these interrelationships, cognitive function and psychopathology symptoms are both important factors to measure and study together in the context of the developing adolescent brain.

An alternative to the statistical methods employed here could be to take a frequentist approach, testing for between-cluster differences in mental health and cognitive function measures by applying analyses of variance or Kruskal-Wallis tests, with post-hoc testing using Tukey’s Honestly Significant Difference or similar (72; 73). When we implemented these methods, we found that the post-hoc analyses were not definitive, and that these methods were underpowered when correcting for multiple comparisons given the small sample size used in the present study. The Bayesian approach used here avoids these barriers and provides more inferentially informative findings, giving direct posterior probabilities for differences in mental health and cognitive function measures between clusters (74).

There are several limitations to the present study. The process of manual EEG feature selection, followed by dimensionality reduction with PCA, faces an obstacle in the number of features we can analyse across a limited number of electrodes, given the small sample sizes typically available in EEG studies. This is because like other popular dimension reduction methods, PCA is known to perform poorly and encounter difficulties as the number of features approaches or exceeds the number of observations — other dimensionality reduction approaches would need to be considered for situations where the number of features of interest exceeds the number of participants for whom data is available. Another limitation of this approach is that it requires manual selection of known or commonly used features calculated from EEG data, which may limit the ability of this approach to find novel patterns or features in EEG signals that are associated with important differences between individuals and subgroups. To improve on these shortcomings, planned extensions to this work include longitudinal analyses of data from a larger number of individuals, and we will investigate alternative approaches to dimension reduction such as functional PCA or manifold learning which may have more flexibility to capture rich information from EEG time series while avoiding some of the challenges around high dimensionality and manual feature selection.

The simple approach implemented here to combined multiple sets of results from different clustering algorithms meant that we could not make inferences about *n* = 11 individuals who are not allocated to final clusters due to being inconsistently clustered across the algorithms). This was suitable given the goal of this work to identify distinct EEG-based subgroups without aiming to support inferences about every individual participant based on their resting state EEG data. An alternative approach in future would be to combine the results using Bayesian model averaging, allowing us to make probabilistic statements about cluster membership and relating this to probability of differences between clusters, taking into account the strength of allocations to the clusters.

Future work in this area will allow us to explore if and how these distinctive patterns of resting state EEG activity between clusters at age 12 can be useful to support longitudinal prediction of cognitive function and mental health outcomes at older ages. Our findings to date suggest potential utility to be explored in developing risk profiles that could be used in a clinical context of risk prediction and early intervention. As more data from LABS become available, our future work will investigate whether the distinctive EEG characteristics of these clusters are useful as an indicator of risk for cognitive development and mental health, both cross-sectionally and over time through adolescent development.

## 6 Acknowledgements

We thank the Longitudinal Adolescent Brain Study (LABS) participants and their caregivers.

